# Chromatin Compaction by Small RNAs and the Nuclear RNAi Machinery in *C. elegans*

**DOI:** 10.1101/469866

**Authors:** Brandon D. Fields, Scott Kennedy

## Abstract

DNA is organized and compacted into higher-order structures in order to fit within nuclei and to facilitate proper gene regulation. Mechanisms by which higher order chromatin structures are established and maintained are poorly understood. In *C. elegans*, nuclear-localized small RNAs engage the nuclear RNAi machinery to regulate gene expression and direct the post-translational modification of histone proteins. Here we confirm a recent report suggesting that nuclear small RNAs are required to initiate or maintain chromatin compaction states in *C. elegans* germ cells. Additionally, we show that experimentally provided small RNAs are sufficient to direct chromatin compaction and that this compaction requires the small RNA-binding Argonaute NRDE-3, the pre-mRNA associated factor NRDE-2, and the HP1-like protein HPL-2. Our results show that small RNAs, acting via the nuclear RNAi machinery and an HP1-like protein, are capable of driving chromatin compaction in *C. elegans*.

## Introduction

Chromatin compaction is necessary for chromosome function. For instance, global chromatin compaction is needed for chromosome segregation during mitosis and meiosis while more localized chromatin compaction is associated with chromosomal structures such as centromeres (constitutive heterochromatin) as well as developmental gene regulation (facultative heterochromatin). Several molecular systems have been identified in eukaryotic cells that mediate chromatin compaction. For instance, the condensin family of proteins hydrolyze ATP to mediate a stepwise compaction of DNA that, in large part, underlies global chromatin compaction preceding mitosis and meiosis ^1,2^. In addition, Polycomb Repressive Complexes (PRCs) 1/2, which regulate HOX genes during development, are two histone-modifying complexes that have been associated with localized chromatin compaction *in vivo* and *in vitro* ^3,4^. Finally, HP1 is a non-histone protein that is a defining feature of heterochromatin in eukaryotes ^5–8^. Tethering HP1 to genes is sufficient to induce chromatin compaction and gene silencing, suggesting that HP1 proteins may directly mediate chromatin compaction and that this compaction may regulate gene expression ^9,10^. HP1 proteins typically possess a chromodomain and a chromoshadow domain ^11,12^. The HP1 chromodomain binds post translationally modified histone 3 such as Histone 3 Lysine 9 trimethylation (H3K9me3), while the chromoshadow domain is important for HP1 homodimerization ^13,14^. HP1-like proteins are able to compact H3K9me3 containing chromatin *in vitro* and the ability of HP1 to mediate chromatin compaction *in vivo* depends upon the ability of HP1 to homodimerize ^15,16^. These observations have led to a model in which HP1-like proteins directly mediate chromatin compaction by bridging distant chromatin sites that harbor H3K9me3.

RNAi is an evolutionarily conserved gene regulatory mechanism triggered by double-stranded RNA (dsRNA). dsRNA is recognized and processed by Dicer-like enzymes into small interfering RNAs (siRNAs) of 21-25 nucleotides in length ^17^. siRNAs are bound by Argonaute (AGO) proteins to form ribonucleoprotein complexes that use the sequence information contained within siRNAs to regulate complementary RNAs *in trans* (via Watson-Crick base pairing) ^18^. In many eukaryotes, siRNAs are found in nuclei where they bind nascent RNAs to co-transcriptionally regulate gene expression as well as direct the deposition of H3K9me3 on chromatin (termed nuclear RNAi) ^18^. In *S. pombe*, nuclear siRNAs, H3K9me3, and HP1 help maintain heterochromatin at the pericentromere ^19^. During this process, an RNA-dependent RNA polymerase enzyme Rdp1 uses nascent RNAs, which are transcribed from the pericentromere, to amplify pericentromere siRNA populations. Amplified siRNAs bind the Argonaute protein Ago1, interact with nascent pericentromeric RNAs, and recruit the H3K9 methyltransferase enzyme Clr4. Once localized, Clr4 generates H3K9me3, which acts as a signal to recruit the HP1-like protein Swi6 as well as promote further Ago1/chromatin interactions ^13,19–25^. Thus, siRNAs, H3K9me3, and HP1 act together in a feed-forward loop to promote heterochromatin formation in fission yeast. Argonaute proteins have been linked linked to H3K9 methylation and heterochromatin formation in many other eukaryotes including, plants, insects, and mammals ^18,26,27^. In some cases, a different class of small RNA, termed piRNA, substitutes for siRNAs during heterochromatin formation ^28^. In summary, an axis of nuclear small RNAs, H3K9 methylases, and HP1-like proteins contribute to heterochromatin formation and chromatin compaction in many eukaryotes.

siRNAs direct H3K9me3 in *C. elegans* ^29^. To do so, cytoplasmic siRNAs engage AGO proteins (HRDE-1 in the germline; NRDE-3 in the soma), which escort siRNAs into nuclei where they bind nascent transcripts (pre-mRNA) and recruit the downstream nuclear RNAi effectors (NRDE-1/2/4) to genomic sites of RNAi ^29–32^. Once recruited, the NRDEs inhibit RNAP II elongation and direct the deposition of H3K9me3 via a currently unknown histone methyltransferase(s) ^29^. *C. elegans* express an abundant class of endogenous (endo) siRNAs, which engage the nuclear RNAi machinery to regulate gene expression and chromatin states (*e.g.* H3K9me3) during the normal course of growth and development ^29–33^. *C. elegans* possess two HP1-like proteins HPL-1 and HPL-2 ^34,35^. HPL-2 is required for RNAi to direct long-term (transgenerational) gene silencing, which is a process known to depend upon the other components of the *C. elegans* nuclear RNAi machinery ^32,36,37^. Thus, HPL-2 may be a downstream component of the nuclear RNAi machinery in *C. elegans*. Surprisingly, the relationship between HPL-2 and H3K9me3 in *C. elegans* is unclear; HPL-2 colocalizes with H3K9me3 on chromatin, however, HPL-2 is still largely able to localize to chromatin (albeit at attenuated levels) in mutant animals that lack detectable levels of H3K9me3 ^38^. Thus, other chromatin marks, in addition to H3K9me3, may contribute to HPL-2 localization in *C. elegans*. In plants, microchidia (MORC) GHKL ATPases are thought to act downstream of siRNAs and H3K9me3 to compact and silence chromatin surrounding transposable elements ^39^. The *C. elegans* genome encodes a single MORC-like protein (MORC-1) ^39^. A recent study showed that MORC-1 is a downstream component of the *C. elegans* nuclear RNAi machinery ^40^. Interestingly, in *C. elegans* lacking MORC-1, germ cell chromatin becomes disorganized and decompacted ^40^). The data suggest that endo siRNAs, acting via the nuclear RNAi machinery, may contribute to chromatin compaction in the *C. elegans* germline.

Here we show that two additional components of the *C. elegans* nuclear RNAi machinery are required for normal chromatin compaction and chromatin organization in the germline, supporting the model that endo siRNAs regulate chromatin compaction in *C. elegans*. In addition, we show that siRNAs are sufficient to direct chromatin compaction in the soma and that this process requires a nuclear RNAi Ago (NRDE-3) as well as the HP1-like factor HPL-2. Our results support a model in which nuclear-localized small regulatory RNAs are important mediators of chromatin organization and compaction in *C. elegans*.

## Results

### Endogenous siRNAs may compact germline chromatin

Small regulatory RNAs, such as siRNAs or piRNAs, have been linked to heterochromatin formation in many eukaryotes, suggesting that small RNAs may act as specificity factors, via Watson-Crick base pairing, for directing heterochromatin formation and chromatin compaction in many eukaryotes ^27^. In addition, the GHKL ATPase MORC-1, a downstream component of the *C. elegans* nuclear RNAi machinery, promotes chromatin organization and chromatin compaction in adult *C. elegans* germ cells, suggesting that siRNAs, acting via the nuclear RNAi machinery and MORC-1, also regulate chromatin compaction in *C. elegans* ^40^. To test this idea, we asked if two additional components of the *C. elegans* nuclear RNAi machinery: the germline expressed nuclear RNAi AGO HRDE-1, and the nuclear RNAi factor NRDE-2, were, like MORC-1, needed for chromatin organization and compaction in adult *C. elegans* germ cells. To do so, we used fluorescence microscopy to visualize a GFP::H2B chromatin marker in wild-type or *hrde-1(-)* and *nrde-2(-)* animals. [Note, it takes 2-3 generations of growth at elevated temperatures (25°C) for chromatin organization defects to be observed in *morc-1(-)* animals ^40^. The reason for this is not known, but may be linked to the role of nuclear RNAi in transgenerational epigenetic gene regulation in the *C. elegans* germline ^32^.] GFP::H2B was monitored over the course of three generations in animals grown at 25°C (Fig. 1A/B). In the first generation of growth at 25°C, germ cell nuclei of wild-type, *hrde-1 (-)*, and *nrde-2(-)* animals appeared similar (Fig. 1A/B). After three generations of growth at 25°C, however, GFP::H2B marked chromatin in *hrde-1 (-)* or *nrde-2(-)* germ cells became disorganized and GFP::H2B signals appeared enlarged (Fig. 1A and Movie 1). We measured the diameter of GFP::H2B fluorescence in randomly chosen nuclei and confirmed that GFP::H2B occupied more space in *hrde-1(-)* or *nrde-2(-)* animals than in wild-type animals after three generations of growth at 25°C (Fig. 1B). Increases in the amount of space occupied by chromatin in *hrde-1(-)* or *nrde-2(-)* animals could be due to decompaction of chromatin or, conceivably, an increase in DNA content. To distinguish between these possibilities, we quantified DAPI fluorescence (see materials and methods) in wild type or *nrde-2(-)* animals after three generations of growth at 25°C. The analysis indicated that wild-type and *nrde-2(-)* nuclei stained with similar amounts of DAPI, suggesting that the increased size of chromatin in *hrde-1(-)* or *nrde-2(-)* nuclei is due to decompaction and not increased DNA content (Fig. S1). The data suggest that germline chromatin becomes disorganized and decompacted in animals lacking the nuclear RNAi factors NRDE-2 and HRDE-1. We used whole chromosome DNA fluorescent *in situ* hybridization (FISH) to test this idea further. We subjected wild-type and *nrde-2(-)* animals, grown at 25°C for three generations, to whole chromosome DNA FISH targeting chromosomes I, II, and III (See Fields et al bioRxiv 2018 for details on chromosome-level DNA FISH). As expected, chromosome I, II, and III DNA FISH stained three distinct regions of subnuclear space, which were concentrated near the nuclear periphery, a pattern consistent with the expected localization of chromosomes in adult pachytene germ cell nuclei (Fig. 1C) ^41^. In *nrde-2(-)* animals, chromosomes I, II, and III DNA FISH signals appeared disorganized and appeared to occupy more space than in wild-type animals (Fig. 1C). We used the Tools for Analysis of Nuclear Genome Organization (TANGO) software to quantify the amount of space occupied by DNA FISH signals in these animals and confirmed that each chromosome occupied more space in *nrde-2(-)* animals than in wild-type animals (Fig. 1D). We conclude that, similar to what has been seen for MORC-1, the nuclear RNAi AGO HRDE-1 and the downstream nuclear RNAi effector NRDE-2 prevent chromatin decompaction in germ cells of adult *C. elegans*.

**Figure 1.**
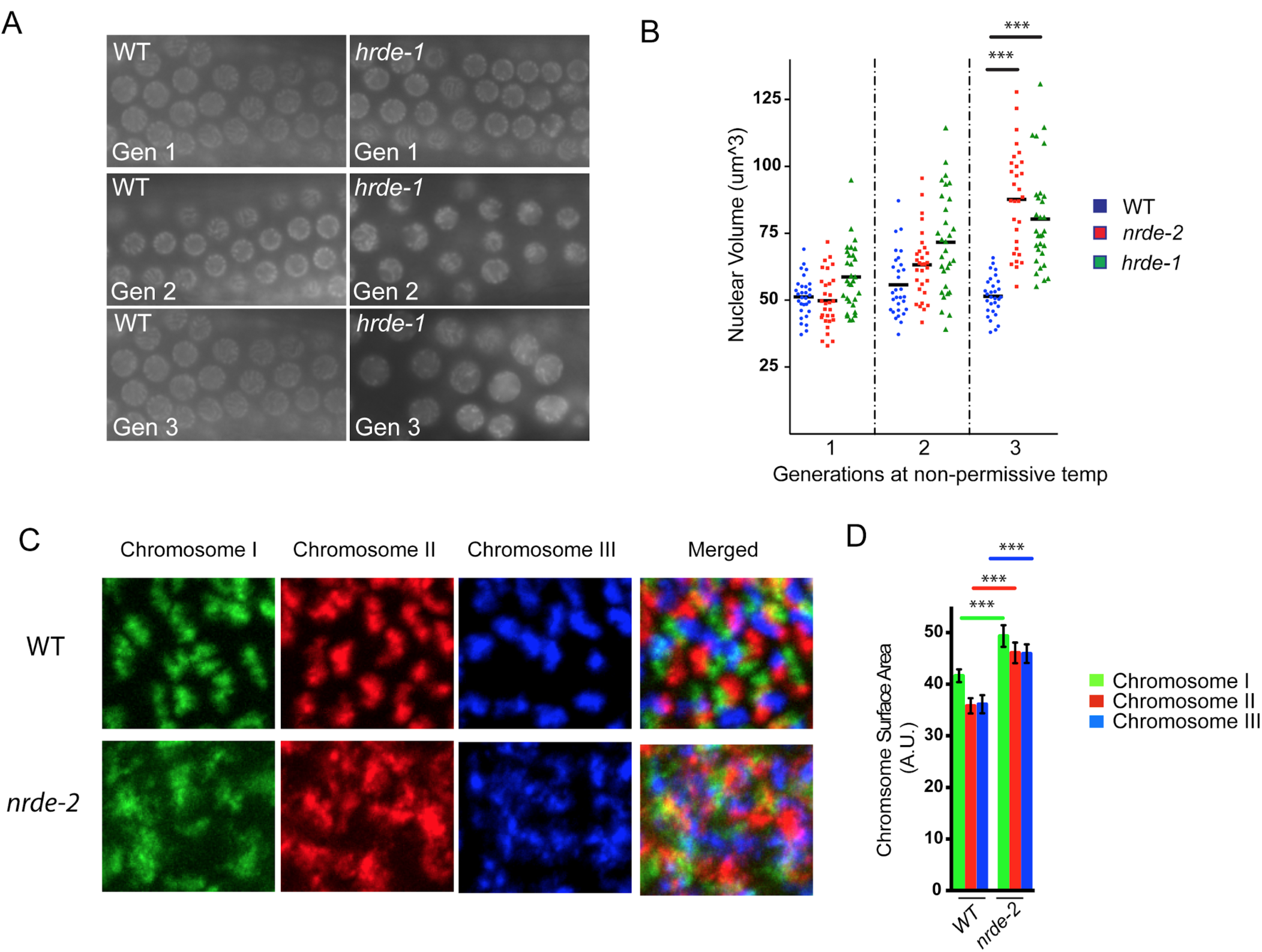
Endogenous siRNAs compact chromatin in *C. elegans.* **A.** Fluorescent micrograph of pachytene germ cells of animals expressing a *gfp::h2b* transgene in the germline as an *in vivo* readout of chromatin structure. Wild type or *hrde-1(tm1200)* animals were maintained at 25°C and imaged over three generations. **B.** Diameter of nuclei in randomly chosen germ cells from panel A were measured (see materials and methods for details) to quantify the nuclear volume of GFP::H2B in wild type, *nrde-2(gg091)*, and *hrde-1 (tm1200)* animals maintained at 25°C across 3 generations. **C.** DNA FISH staining of chromosomes I, II and III in pachthyne germ cells of wild type and *nrde-2(gg091)* animals that were maintained at 25°C for 3 generations. **D.** TANGO-based quantification of randomly chosen germ cells from panel C (see materials and methods for details) of chromosomal surface areas for wild type and *nrde-2(gg091)* animals maintained at 25°C for 3 generations. p-values were calculated using a student’s two-tailed T test. *** = p-value <0.005

### RNAi induces chromatin compaction in *C. elegans*

Our data are consistent with the idea that endogenously expressed siRNAs promote chromatin organization and compaction in *C. elegans*. It is also possible that chromatin decompaction in *hrde-1(-)*, *nrde-2(-),* or *morc-1(-)* animals could be an indirect consequence of the gene misregulation that occurs in animals lacking a functioning nuclear RNAi system ^32,42^. To ask if small RNAs directly mediate chromatin compaction in *C. elegans*, we asked if experimental RNAi were sufficient to drive chromatin compaction. We targeted a large and repetitive multi-copy *sur-5::gfp* transgene, which expresses GFP in all somatic cells of *C. elegans*, with *gfp* RNAi and used *gfp* DNA FISH to visualize the space occupied by *gfp* DNA before and after *gfp* RNAi (Fig. 2A) ^43,44^. The repetitive *sur-5::gfp* transgene was chosen as its large size might be expected to make quantifications of size feasible. Similarly, we chose to image DNA FISH signals in intestinal cells as these cells are polyploid (32N), large, and easy to identify ^45^. After *gfp* RNAi, the subnuclear space occupied by *sur-5::gfp* DNA appeared to decrease (Fig. 2A). Quantification confirmed that *gfp* RNAi caused *gfp* DNA FISH signals to occupy ∼2 fold less space than in animals not exposed to *gfp* RNAi (Fig. 2B and Fig. S2). We conclude that RNAi can induce chromatin compaction in *C. elegans*.

**Figure 2.**
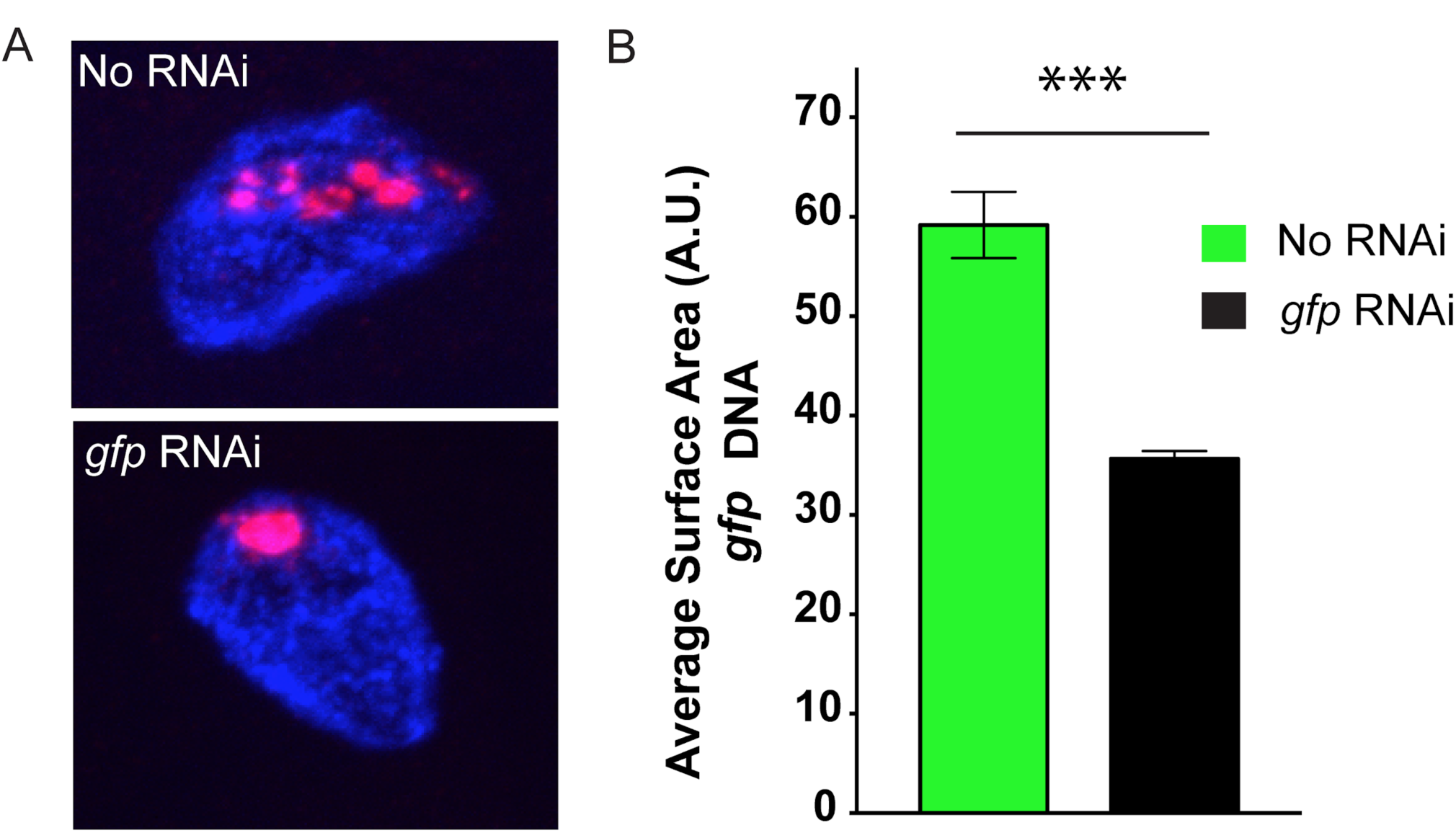
dsRNA induces chromatin compaction in *C. elegans*. **A.** Fluorescent micrographs of a single *C. elegans* intestinal nucleus possessing a multicopy *sur-5::gfp* transgene stained with DNA FISH probes targeting *gfp* DNA (red), and DAPI (blue). Representative images of animals grown in the presence (bottom) or absence (top) of *gfp* dsRNA. **B.** Surface area calculations for *gfp* DNA FISH signals in animals grown in the presence (right) or absence (left) of *gfp* dsRNA. p-value was calculated using a student’s two tailed t-test. *** = p-value <0.005

### Nuclear RNAi couples siRNAs to chromatin compaction

Nuclear RNAi in somatic tissues requires the the somatically expressed AGO NRDE-3 and the conserved pre-mRNA binding protein NRDE-2 ^29,30^. To explore how RNAi mediates chromatin compaction, we asked if *nrde-2(-)* or *nrde-3(-)* animals were able to compact *sur-5::gfp* chromatin in response to *gfp* RNAi. We conducted *gfp* DNA FISH on *nrde-2(-);sur-5::gfp* and *nrde-3(-);sur-5::gfp* animals that were treated +/- with *gfp* RNAi and found that both NRDE-2 and NRDE-3 were required for RNAi-directed chromatin compaction (Fig. 3A/B). Given that NRDE-3 is an AGO, the data support the idea that small RNAs are needed to mediate *sur-5::gfp* chromatin compaction. Given that NRDE-2 is a nuclear RNAi factor, the data support the idea that small RNAs engage the nuclear RNAi machinery to mediate chromatin compaction. In the absence of *gfp* RNAi the space occupied by the *sur-5::gfp* transgene was greater in *nrde-2(-)* and *nrde-3(-)* than in wild-type animals, suggesting that small RNA and nuclear RNAi-based *sur-5::gfp* compaction occurs at some level even in the absence of exogenous sources of *gfp* dsRNA (Fig. 3B). This latter observation is consistent with previous reports demonstrating that repetitive transgenes are often subjected to RNAi-based gene silencing in *C. elegans* ^46^. How might siRNAs direct chromatin compaction? In many eukaryotes including *C. elegans*, RNAi directs H3K9me3 at genomic loci exhibiting sequence homology to RNAi triggers ^27^. In addition, HP1-like proteins are recruited to H3K9me3 containing chromatin and homo-dimerize (or undergo phase separation) to compact chromatin by linking together distant sites of chromatin ^13,16,47^. Finally, the HP1-like protein HPL-2 has been functionally linked to nuclear RNAi in *C. elegans* ^37^. For all these reasons, we asked if HPL-2 might be required for RNAi-mediated chromatin compaction in *C. elegans*. *hpl-2(-)* animals failed to compact *sur-5::gfp* chromatin in response to *gfp* RNAi (Fig. 3B). The data are consistent with the idea that HP1-like proteins contribute to chromatin compaction directed by nuclear siRNAs in *C. elegans*.

**Figure 3.**
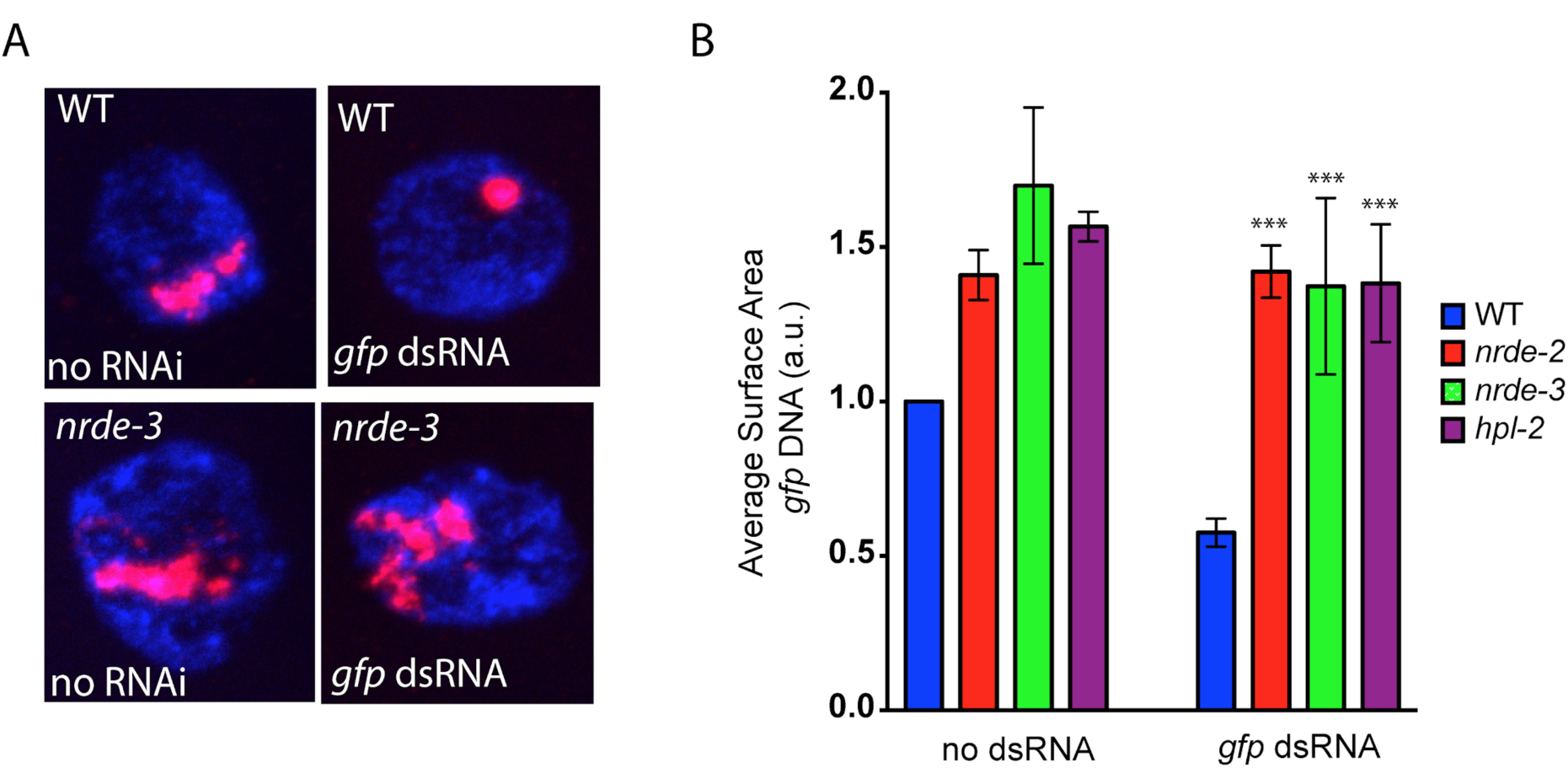
Nuclear RNAi and HPL-2 are needed for RNAi-directed chromatin compaction. **A.** Fluorescent micrographs of intestinal nuclei from animals possessing a multicopy *sur-5::gfp* transgene stained with DNA FISH probes targeting *gfp* DNA (red) and DAPI (blue) in wild type or *nrde-3(gg066)* mutant animals exposed to no RNAi (left) or *gfp* RNAi (right). **B.** Surface area calculations of *gfp* DNA of intestinal nuclei from wild type, *nrde-2(gg091)*, *nrde-3(gg066)*, and *hpl-2(tm1489)* animals, which all express the *sur-5p::gfp* transgene, and which were grown in the presence or absence of *gfp* RNAi. Data is normalized to the WT (no RNAi) control for each condition to account for differences in hybridization efficiency across experiments. p-values were calculated using a student’s two tailed t-test and were calculated with respect to wild type samples feeding on *gfp* RNAi. ***=p-value <0.005. n.s.=not significant.

## Discussion

Here we present evidence supporting the idea that endogenous siRNAs and the nuclear RNAi machinery can initiate/maintain chromatin architecture and compaction in *C. elegans* germ cells. We also show that experimentally introduced siRNAs are sufficient to compact chromatin in the soma and that this chromatin compaction requires a nuclear AGO as well as a downstream component of the nuclear RNAi machinery and the HP1-like protein HPL-2. The data support a model whereby nuclear small RNAs are important regulators of chromatin organization and chromatin compaction in *C. elegans*.

Our genetic analyses identified three factors (NRDE-2, NRDE-3, and HPL-2) that are required for RNAi-directed chromatin compaction in intestinal cells. These results have led us to propose the model summarized in Fig. 4. The AGOs NRDE-3 (soma) and HRDE-1 (germline) bind siRNAs and interacts with homologous nascent RNAs to recruit the NRDE nuclear RNAi factors to chromatin. The downstream nuclear RNAi factors (e.g. NRDE-2) recruit a currently unknown histone H3K9 methyltransferase(s) to chromatin to deposit H3K9me3. Finally, H3K9me3 recruits the HP1-like protein HPL-2, which dimerizes to compact the chromatin (Fig. 4). Important tests of this model will include 1) asking if siRNAs are sufficient to direct chromatin compaction in germ cells, 2) asking if HPL-2 is physically recruited to sites of nuclear RNAi, 3) asking if HPL-2 dimerization is necessary for RNAi-based chromatin compaction, and 4) asking if H3K9me3 (directed by nuclear RNAi) is needed for recruitment of HPL-2 to chromatin. Regarding this latter question, three SET domain proteins (SET-25, MET-2, and SET-32/HRDE-3) have been linked to H3K9 methylation in *C. elegans* and are redundantly required for RNAi directed H3K9me3 ^48–50^. Potential roles for H3K9me3 in HPL-2 recruitment and/ or chromatin compaction could be addressed by asking if animals lacking all three of methytransferases are defective for small RNA-directed chromatin compaction. Interestingly, in *C. elegans*, localization of HPL-2 to chromatin is only partially dependent upon H3K9me3, suggesting that *C. elegans* may possess H3K9me3-independent systems for recruiting HPL-2 to chromatin ^38^. Indeed, *in vitro* studies show that *C. elegans* HPL-2 binds H3K27me3 (as well as H3K9me3) and *in vivo* studies show that RNAi directs the deposition of H3K27me3 on chromatin (as well as H3K9me3) in *C. elegans* ^51,52^. Thus, H3K27me3 may act in parallel with, or independently of, H3K9me3 during HPL-2 driven chromatin compaction. Studies asking if components of the *C. elegans* Polymcomb 2 complex are needed for RNAi-directed chromatin compaction could test this model. Finally, the GHKL ATPase MORC-1 has been linked to chromatin compaction in plants and in *C. elegans* germ cells ^39,40^. These observations have led to a model in which MORC-1 compacts chromatin at the direction of small RNAs. The chromatin compaction system described here will allow the role of MORC-1 in somatic chromatin compaction to be assessed.

**Figure 4.**
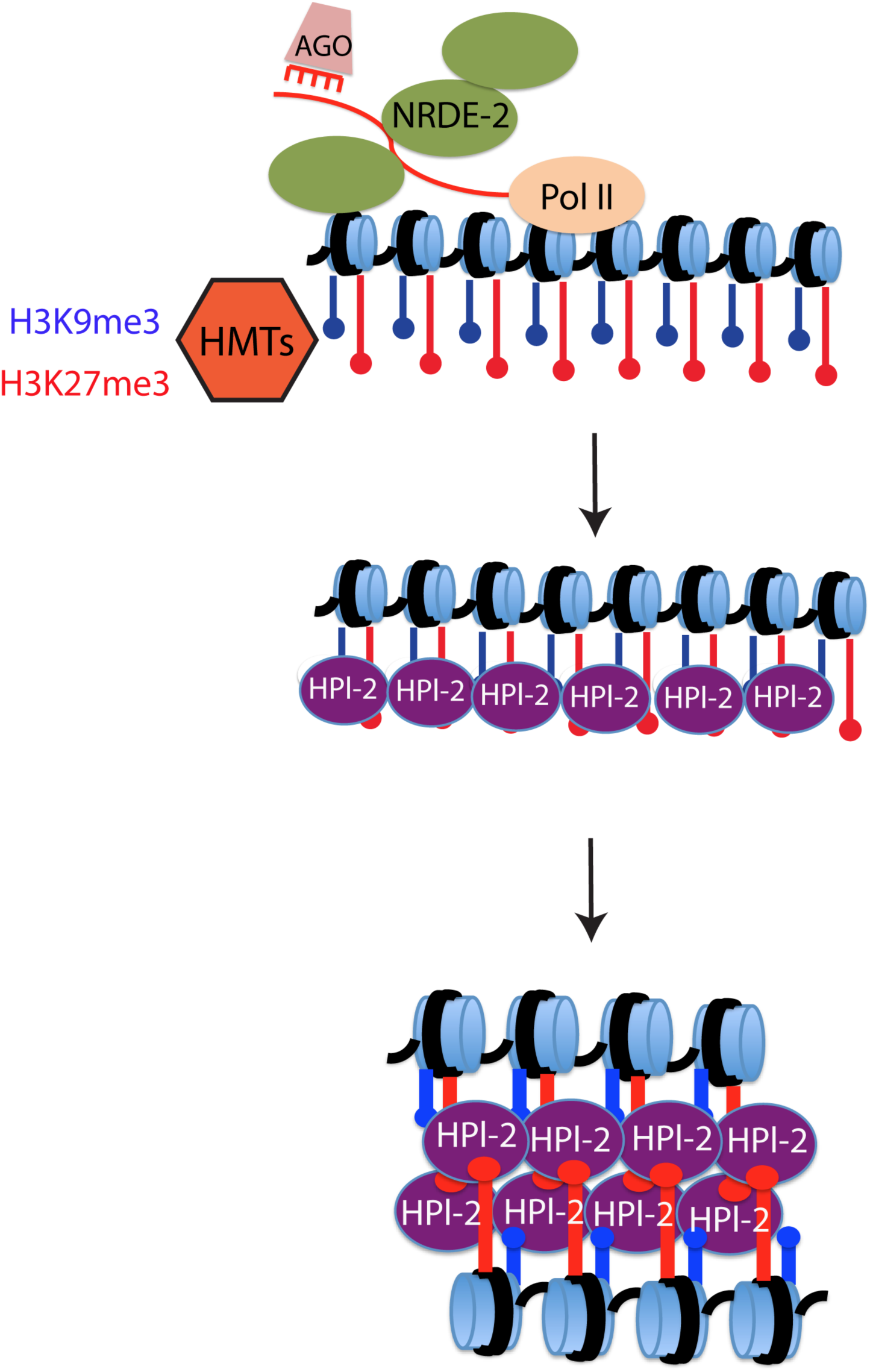
Model for small RNA directed chromatin compaction in *C. elegans.*

Many eukaryotes express a diverse array of small regulatory RNAs. These RNAs are mobile and have the ability to interact with a high degree of specifically with other cellular nucleic acids. Such traits make small regulatory RNAs excellent candidates to be precise yet versatile vectors of chromatin compaction during reproduction and development. What biological function(s) might small RNA-directed chromatin compaction play during reproduction and development? The simplest answer might be that small RNA-directed chromatin compaction regulates gene expression programs by regulating accessibility of chromatin to the transcriptional machinery. Consistent with this idea, the nuclear RNAi factors that we have linked to chromatin compaction in this work, are known to regulate gene expression in *C. elegans* ^29,30,32,42^. Asking if nuclear RNAi-regulated genes are bound by HPL-2 (and if this binding requires nuclear RNAi factors) would be a good test of this idea. Interestingly, many of the loci regulated by nuclear RNAi are not protein-coding genes; rather, they are cryptic loci or pseudogenes ^32,42^. Thus, gene regulation may not be the sole biological output for small RNA-mediated chromatin compaction. It is possible that small RNA- directed chromatin compaction could also be used to help generate higher-order chromatin structures underlying chromosome segregation during mitosis or meiosis. Consistent with this latter idea, mutations in several *C. elegans* RNAi factors (including HRDE-1 and NRDE-2) have been linked to chromosomal nondisjunction during meiosis ^53–62^. Centromeres are compacted chromosomal structures needed for chromosome segregation during mitosis and meiosis. *C. elegans* are holocentric; centromere function is dispersed across hundreds of point centromeres on each chromosome ^63^. Nonetheless, like centromeres in other eukaryotes, *C. elegans* point centromeres are flanked by chromatin containing H3K9me3 and HP1 (termed pericentromeres) and this flanking heterochromatin is important for centromere function ^38,64,65^. Therefore, it is possible that small RNA-directed HPL-2 recruitment and, therefore, chromatin compaction, may contribute to holocentromere function in *C. elegans*, an idea that could be tested by asking if the localization of HPL-2 to pericentromeres depends upon nuclear small RNAs.

**Supplementary Figure 1.**
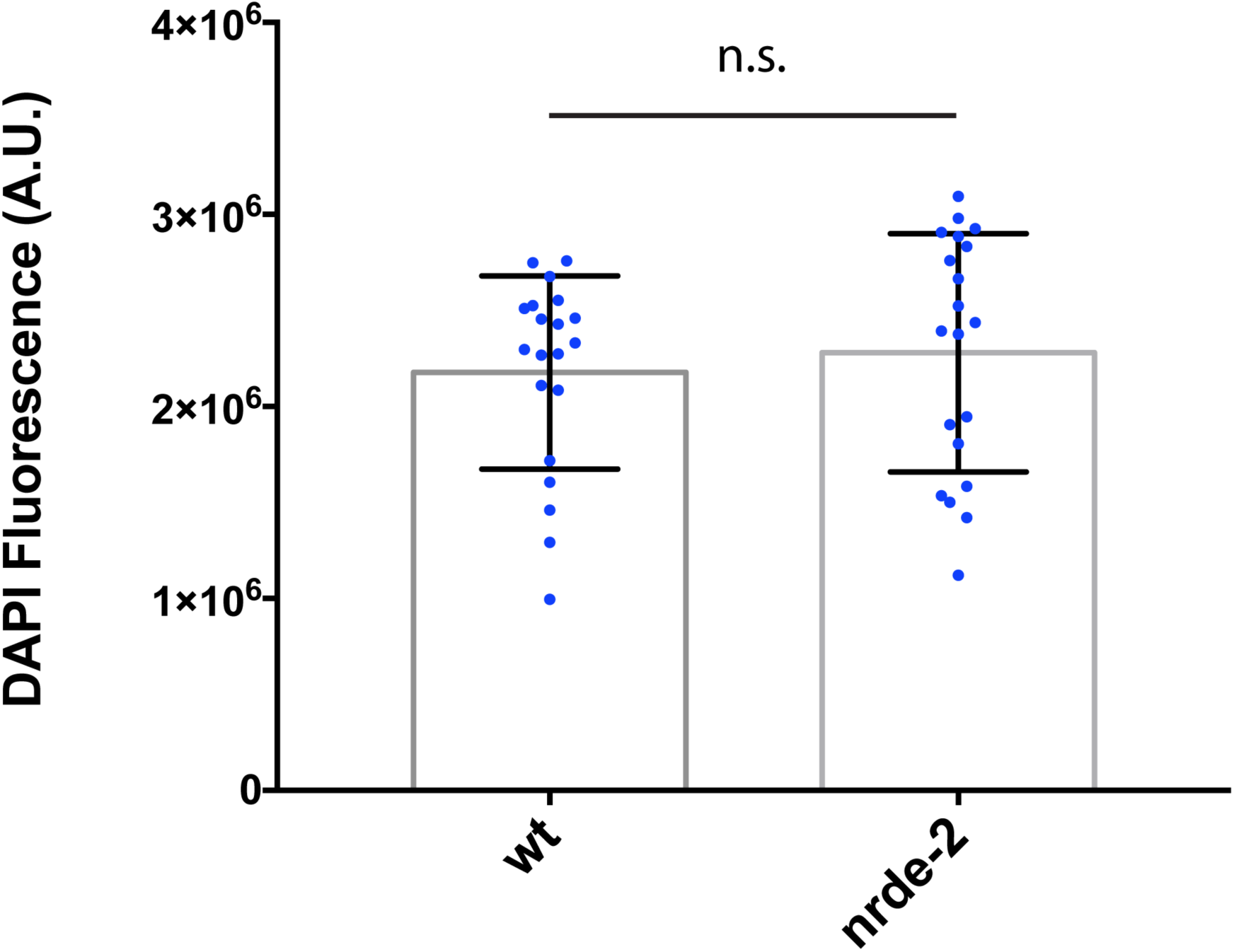
Wild-type and late generation *nrde-2(-)* nuclei possess a similar amount of DNA. Fluorescent intensity measurements were made from individual germ cells of wild type and *nrde-2(gg091)* animals grown at 25°C for three generations. Each data point represents one germ cell nuclei. Data points were collected from four different animals per genotype. p-value was calculated using a student’s two tailed t-test. n.s. = not significant.

**Supplementary Figure 2.**
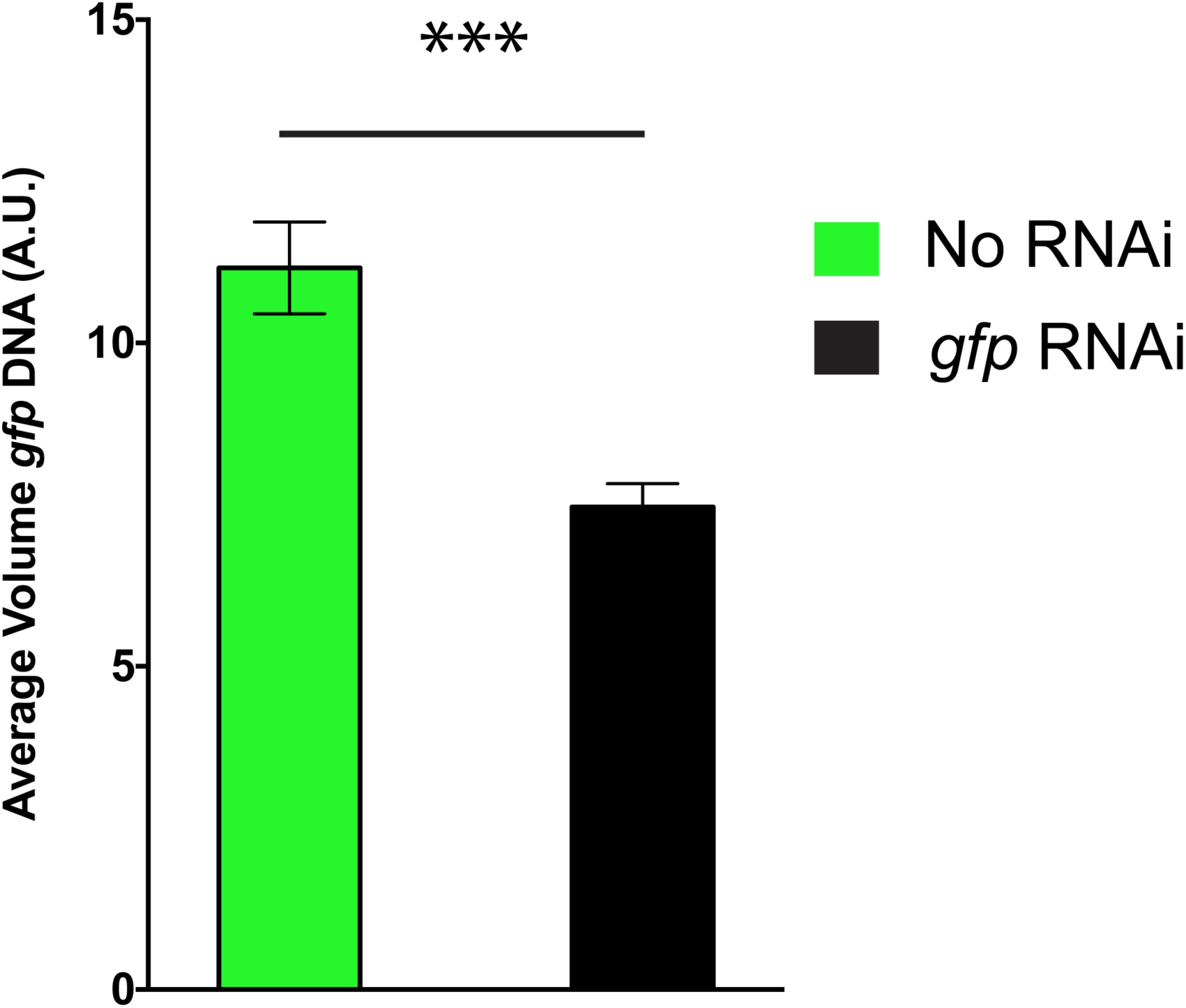
Volume quantifications of the *sur-5::gfp* transgene. Volume quantifications for animals feeding on *gfp* RNAi (right) or no RNAi (left). p-value was calculated using a student’s two tailed t-test. *** = p-value <0.005.

**Movie 1. 3D reconstructions of GFP::H2B in wild type versus *nrde-2(-)* mutant animals.** 3D reconstructions of confocal Z-slices for wild type and *nrde-2(gg091)* animals expressing a *gfp::h2b* reporter in the germline and maintained at 25°C for three generations. Z-slices were gathered at 0.3 um intervals. 3D reconstructions were generated using Nikon Imaging software.

## Materials and Methods

### Strains

N2, MH1870 (kuIs54), YY519 (*nrde-2(gg091); kuIs54*), YY520 (*nrde-3(gg066); kuIs54*), YY528 *(hrde-1 (tm1220); pkIs32)*, YY1704 (*nrde-2 (gg091); pkIs32*), YY502 (nrde-2(*gg091*)), YY1363 (*hpl-2 (tm1489), kuIs54*).

### Sample collection for DNA FISH

Embryos were isolated by hypochlorite treatment and placed on 10 cm plates seeded with OP50 bacteria, or for RNAi experiments HT115 bacteria expressing *gfp* dsRNA or no dsRNA. When animals reached adulthood, plates were washed with M9 solution and collected in 15 ml conical tubes. Animals were pelleted (3k rpm for 30 seconds), and washed 2 times with M9 solution. Animals were resuspended in 10 ml of M9 solution and rocked for ∼30 min at room temperature. Animals were pelleted and aliquoted to 1.5 ml microcentrifuge tubes (40 ul of packed worms per tube). Samples were placed in liquid nitrogen for 1 minute and stored at −80C.

### DNA FISH probe preparation

Genomic DNA was isolated from animals expressing *kuIs54* (*Psur-5::sur-5::gfp*). PCR was done to amplify the *gfp* sequence, and 2 ug of *gfp* DNA was used as starting material for FISH probe synthesis. The FISH Tag DNA Kit from Life Science Technologies (cat. number F32948) was used and the protocol suggested by the manufacturer was followed. Exceptions to the manufacturer's protocol were the amount of starting material (2 ug used) and resuspension volume (20 ul). Probes were stored at −20C in the dark and used within 2 weeks of synthesis.

### *gfp* DNA FISH

Frozen worm pellets were resuspended in cold 95% ethanol and vortexed for 30 seconds. Samples were rocked for 10 minutes at room temperature. Samples were spun down (3k rpm for 30 seconds) and supernatant discarded. Samples were washed twice in 1X PBST. 1 ml of 4% paraformaldehyde solution was added and samples were rocked at room temperature for 5 minutes. Samples were then washed twice with 1X PBST before resuspension in 2XSSC for 5 minutes at room temperature. Samples were spun down and resuspended in a 50% formamide 2XSSC solution at room temperature for 5 minutes, 95°C for 3 minutes, and 60°C for 20 minutes. Samples were spun and resuspended in 60 ul of hybridization mixture (10% dextran sulfate, 2XSSC, 50% formamide, 2 ul of *gfp* FISH probe and 2 ul of RNAse A (sigma 20 mg/ml)). Hybridization reactions were incubated at 95°C for 5 minutes before overnight incubation at 37°C in a hybridization oven. The next day, samples were washed with 50% formamide 2XSSCT (rotating at 37°C) for 30 minutes. Wash buffer was removed and samples were resuspended in mounting medium (vectashield with DAPI). Samples were mounted on microscope slides and sealed with nail polish.

### Oligopaint DNA FISH

For Oligopaint staining, the same DNA FISH protocol was used with the following exceptions. In the hybridization mixture, 100 pmol of oligo was used for each chromosome. The following day, samples were washed with 50% formamide 2XSSCT at 37C for 30 minutes. Wash buffer was removed, and samples were resuspended in in 60 ul of hybridization mixture (10% dextran sulfate, 2XSSC, 50% formamide, and 100 pmol of bridge oligo for each chromosome). Samples were incubated at 37C for 45 minutes. Samples were washed with 50% formamide 2XSSCT at 37C for 30 minutes. Wash buffer was removed, and samples were resuspended in 60 ul of hybridization mixture (10% dextran sulfate, 2XSSC, 50% formamide, and 100 pmol of detection oligo (labeled with Alexa 488, cy3, or Alexa647) for each chromosome). Samples were incubated at 37C for 45 minutes. Samples were washed with 50% formamide 2XSSCT at 37C for 30 minutes. Wash buffer was removed, and samples were resuspended in 50 ul of slowfade gold with DAPI.

### DAPI staining alone

Frozen worm pellets were resuspended in 1 ml of cold methanol (−20C). Samples were rocked for 15 minutes at room temperature. Samples were washed twice with 1XPBST, and resuspended in 75 ul of vectashield with DAPI.

### Microscopy

DNA FISH images were captured by standard fluorescent microscopy using a widefield Zeiss Axio Observer.Z1 microscope using a Plan-Apochromat 63X/1.40 Oil DIC M27 objective and an ORCA-Flash 4.0 CMOS Camera. The Zeiss Apotome 2.0 was used for structured illumination microscopy using 3 phase images for DNA FISH imaging. For each animal imaged, the exposure time was set to fill 35% of the camera's dynamic range to account for differences in hybridization efficiency between animals. For intestinal nuclei, all images were taken at the anterior end of the worm to ensure we were imaging the same set of nuclei across all images. All image processing was done using the Zen imaging software from Zeiss. For *in vivo* imaging of GFP::H2B, animals were immobilized in M9 with 0.1% Sodium Azide. Animals were imaged immediately with the Zeiss Axio system described above. Confocal imaging (Movie 1) was done using a Nikon Eclipse Ti microscope equipped with a W1 Yokogawa Spinning disk with 50 um pinhole disk and an Andor Zyla 4.2 Plus sCMOS monochrome camera. A 60X/1.4 Plan Apo Oil objective was used.

### Quantification of space occupied by GFP::H2B

To measure volume of H2B::GFP signals in Fig. 1B, nuclear diameters were measured using the Zeiss Zen software on randomly selected nuclei located in a similar region of the germline (close to gonad turn, pachytene). Nuclear volumes were calculated assuming spherical shape.

### Quantification of space occupied by DNA FISH signals

The Tools for Analysis of Nuclear Genome Organization (TANGO) software was used to quantify the surface area and/or volume of space occupied by DNA FISH signals (*gfp* DNA FISH and whole chromosome DNA FISH ^66^. For *gfp* DNA FISH analysis in intestinal nuclei, the standard nucleus segmentation processing chain was used with the following modifications: Nucleus edge detector, minimum nucleus size = 15,000. Size and edge filter, minimum nucleus size = 15,000. For object (*gfp* DNA FISH signal) segmentation the nucleoli processing chain was used with the following modification: Size and edge filter, minimum volume = 3. For whole chromosome DNA FISH quantifications, the standard nucleus segmentation processing strain was used with the following modifications: Nucleus edge detector, minimum nucleus size = 100. Size and edge filter, minimum nucleus size = 100. The nucleoli processing chain was used to segment chromosome structures with the following modifications: Size and edge filter, minimum volume = 3.

### DAPI quantifications

As a readout of DNA quantity we quantified DAPI fluorescence using ImageJ. Regions of interest (ROI), were generated for five germ cell nuclei per worm, and z-slices were selected to encompass the entire nucleus. The multi-measure tool in ImageJ was used to measure total fluorescence within the ROI for each slice. Three ROIs were drawn outside the worm to gain an average background level. Background levels were subtracted from each nucleus by multiplying the area of the ROI for a given nucleus with the average background value for the image.

## Data availability

All data generated or analysed during this study are included in this published article (and its Supplementary Information files).

## Author Contributions

B.D.F generated all data. B.D.F and S.K. wrote the paper

